# Differential participation of the corticospinal and corticorubral neurons during motor execution in the rat

**DOI:** 10.1101/2024.09.24.614822

**Authors:** Paola Rodriguez-Moreno, Juliana Loza-Vaqueiro, Rafael Olivares-Moreno, Gerardo Rojas-Piloni

## Abstract

The sensorimotor cortex is crucial for learning and executing new movements with precision (Nudo & Frost, 2007). It selectively modulates sensory information flow and represents motor information in a spatially organized manner (Canedo, 1997; Chen et al., 2017). The pyramidal system is made up layer 5 pyramidal tract neurons (PTNs), which are organized into populations with distinct morphological, genetic and functional properties. These subpopulations project to different subcortical structures in a segregated manner (Nudo & Frost, 2007).

To understand whether PTNs projecting to different structures play distinct functional roles in motor control, we characterized two types of layer 5 neurons in the motor cortex: corticorubral (CR) neurons, which project to the red nucleus, and corticospinal (CS) neurons, which project to the spinal cord.

To analyze movement performance in rats, we compared the selective optogenetic inhibition of motor cortex CS or CR neurons during lever movement execution in response to a light stimulus. As the animals progressed through the training sessions, the variability of lever trajectories decreased, and the movements became more stereotyped. Photoinhibition of CS or CR neurons increased the performance variability of learned movements but differentially affected kinematic parameters. CR neuron inhibition affected amplitude, duration, reaction times, speed, and acceleration of the movement. In contrast, inhibition of CS neurons mainly altered the duration, speed, and acceleration of the movement. We conclude that CS and CR are complementary pathways for transmitting information rather than copies of the same motor command.

## INTRODUCTION

As the main cortical output, the PTNs receives and integrates inputs to coordinate the activity of numerous subcortical structures involved in movement execution, including the striatum (corticostriatal pathways), the pontine nucleus (corticopontine pathway), the spinal cord (corticospinal [CS] pathway), and the red nucleus (corticorubral [CR] pathway) (Isomura et al., 2009; Li et al., 2015).

Research on CR projections has been limited. From an evolutionary perspective, animals with undulating locomotion mechanisms lack the magnocellular zone of the red nucleus and the rubrospinal tract, which appear early in the radiation of mandibulated vertebrates as a limb motor control system (Nudo & Frost, 2007; Olivares-Moreno et al., 2021). The rubrospinal tract originates mainly from the ventromedial part of the magnocellular region of the red nucleus (Liang et al., 2012; Massion, 1967), although some authors have shown that the parvocellular region also contributes to this tract in rodents and cats (Huisman et al., 1981, 1983; Shieh et al., 1983). The rubrospinal tract ends in the lateral part of the intermediate zone (Rexed laminae V to VII) at all levels of the spinal cord (Strominger et al., 1987), contacting with dendritic branches of premotor interneurons (Kostyuk & Skibo, 1975).

The parvocellular region of the red nucleus has projections to the ipsilateral inferior olive, which in turn projects to the cerebellum (Edwards, 1972; Kennedy & Humphrey, 1987; Massion, 1967; Walberg & Nordby, 1981). On the other hand, the parvocellular zone receives afferents from the dentate nucleus of the cerebellum (Angaut & David, 1965; Courville, 1966) and is innervated directly by the cerebral cortex through the CR pathway and indirectly by collaterals of the pyramidal tract and CS neurons (Brown, 1974; Canedo, 1997; Gwyn & Flumerfelt, 1974; Tsukahara et al., 1967, 1968).

The CS tract arises evolutionarily in parallel with the development of the parvocellular zone of the red nucleus (Canedo, 1997, 2003). Although the CS tract plays a fundamental role in motor control, it is a complex multifunctional pathway composed of subsystems that selectively modulate sensory information in the dorsal horn by depolarizing primary afferents (Lemon, 2008). It also participates in sensorimotor integration by modulating motor execution directly through projections to ventral horn motoneurons and indirectly through premotor interneurons that influence spinal cord motoneurons (Moreno-López et al., 2016; Olivares-Moreno et al., 2017).

Previously, has been documented CS and rubrospinal tract interaction during lesion experiments. In this sense, CS tract function could compensate for motor deficits caused by damage to the red nucleus or loss of the corticorubrospinal pathway. Similarly, plasticity has been reported in the corticorubrospinal tract when there is damage to the CS tract (Humphrey & Rietz, 1976; Williams et al., 2014). The fact that CS and corticorubrospinal projections converge on their objectives within the spinal cord, modulating local interneurons and propriospinal neurons (Alstermark et al., 1981; Illert et al., 1976, 1977), allow to suggest that there is a close relationship between these two motor pathways.

Here we show that CS and CR are two PTNs populations located in layer 5B of the sensorimotor cortices. In fact, PTNs are organized into populations that exhibit differential morphological, genetic, hodological, and functional properties (Economo et al., 2018; Rojas-Piloni et al., 2017). The segregated subcortical projections of these neurons (Economo et al., 2018; Rojas-Piloni et al., 2017) indicate that the sensorimotor cortex is capable of processing different types of information and producing individual output channels to subcortical structures, thereby modulating various functions associated with sensorimotor control. However, little is known about the organization of subcortical projections, intracortical microcircuitry, and synaptic interactions in the sensorimotor cortex that encode cortical outputs. In this study, we provide new evidence on the organization and special characteristics of two pyramidal system pathways. Our results demonstrate that the CR tract is a valuable object of study, since the information on its functional role within sensorimotor control in physiological conditions is scarce and poorly updated.

## MATERIAL AND METHODS

### Animals

Twenty-one-day-old Wistar rats were used in this study. They were housed under controlled temperature (23 ± 2°C) and light conditions (12-hour light/dark cycle) with *ad libitum* access to a standard laboratory diet (Lab Diet-5001). The animals were handled daily for 3 days prior to the behavioral task to minimize stress. All experimental procedures were conducted in accordance with the guidelines of the Bioethics Committee of the Institute of Neurobiology (protocol No. 073) and the Official Mexican Standard NOM-062-ZOO-1999.

### Behavior

Water was restricted for 24 hours a day, 5 days a week. The animals received water only during the training sessions and one hour after. The task was conducted in black polycarbonate boxes (39 cm wide x 47 cm long x 20 cm high) in a completely dark room. Each box contained a light source on the back wall, a standard lever (Lafayette Instrument model 80110) positioned 1.5 cm from the floor, and a licking port 6 cm from the floor connected to a peristaltic pump (Lafayette Instrument model 80204P). The Animal Behavior Environment Test program (ABET II Standard, Lafayette Instrument version 2.16.1) automatically controlled the experiment.

Digital light and analog lever signals were recorded using the Open Ephys program (Open Ephys Acquisition Board ®) at a sampling frequency of 5000 Hz. Additionally, infrared sensors tracked the vertical movement of the lever during the behavioral task.

The task consisted of a lever press movement with the left forelimb in response to a light stimulus. For each correct trial execution, the animal received a reinforcing stimulus of 15 μL of water. During training, the light stimulus or delay varied from durations of 700 ms to 1300 ms during the sessions. A 2000 ms response time started after the light was turned off (Go signal). During this time, the rats had to press the lever to receive the reward. Afterwards, an inter-trial time began, ranging randomly from 2000 ms to 4000 ms (Figure 2 A).

A trial was considered correct when the animal executed the lever movement within the response time. An incorrect trial was categorized as “anticipated,” if the animal performed the lever movement during the light stimulus, or “out of time,” if the response occurred between trials. Incorrect trials were penalized with a timeout (random time between 2000 ms and 5000 ms) before initiating a new trial. Finally, an omission was defined as the failure to respond to the light stimulus and execute the lever movement.

### Surgeries

#### Injections

For the retrograde viral or tracer injections, rats were anesthetized with isoflurane in O2 (5% to induce and 1.2% to maintain) (Sofloran® Vet) and placed in a stereotaxic apparatus. Local anesthesia (Piscaine 2%) was injected subcutaneously into the skull. To label CS neurons, the retrograde virus was injected (120 nL) between the C4-C5 and C5-C6 intervertebral space on the left side of the spinal cord (ML:1.00). For the CR neurons, a trephine was made with a dental drill at the coordinates of the red nucleus (AP: 5.7; ML: 1.00), followed by the injection of the retrograde virus directly into the red nucleus (20 nL DV: 6.8). Injections were performed with graduated pipettes with a tip diameter of 10 to 25 µm (BlauBrand intraMARK ref 7087 07), coupled to a pneumatic picopump (PV830, WPI), enabling discrete injections (10 to 30 nL) with pressure pulses of 100 ms to 20 PSI. After injection, the pipette was left at the injection site for 3 minutes before removal. Wounds and muscles were closed with absorbable sutures (Atramat ® Silk USP 3-0 and 6-0) and liquid suture (Vetbond – Tissue Adhesive). For pain control, TRAMADOL® (25 mg/kg) was administered every 12 hours, and Metamizole Sodium (0.1 ml/100 g) was given every 12 hours for 3 post-surgical days.

The injected retrograde viruses were pAAV-hsyn-Jaws-KGC-GFP-ER2 (Addgene; Plasmid #65014) and AVV-Syn_ChR2(H134R)-GFP (control virus; Addgene; Plasmid #58880). The retrograde tracers injected were Fluorogold 2% (Fluorochrome) into the red nucleus and BDA 0.1% (Invitrogen, 3000 MW, Anionic, ref D3307) into the spinal cord.

#### Optical fiber implantation

Four weeks after viral injections, the rats were anesthetized with isoflurane (Sofloran® Vet) and placed in a stereotaxic apparatus. A trephine was performed with a dental drill at the coordinates of the M1 cortex (AP: 2.20; ML: 2.00; DV: 1.00; Supplementary Fig. 3). The optical fiber (Doric lenses® 1 mm long) was implanted and cemented (Ortho-Jet Power).

### Optogenetic inhibition through photostimulation

The optogenetic inhibition of CS (n=4) and CR (n=4) neurons was performed by means of light pulses (620 nm; 90 mW/mm^2^) controlled by Radiant® software (PlexBright® Optogenetic Stimulation System). Photoinhibition was randomly applied in two distinct phases of the learned motor task, either concurrently with the light stimulus (i.e., early inhibition) for 1300 ms or after the light stimulus (i.e., late inhibition) for 2000 ms (Fig. 3D). During the experiment, each 60-minute session consisted in an equivalent number of trials with optogenetic inhibition and without photostimulation (control phase).

### Histology (image acquisition and analysis)

Following the optogenetic inhibition sessions through photostimulation, rats were deeply anesthetized with pentobarbital (PISABENTAL®; 45 mg/kg i.p.) and intracardially perfused with 250 ml of 0.1 M PB (phosphate buffer) followed by 250 ml of PFA (paraformaldehyde). The spinal cords and brains were stored in PFA at 4° C for three days. Coronal slices (70 µm thickness) were made on a vibratome (Leica® VT1200 S). Subsequently, the sections were mounted on slides for image acquisition using a fluorescence microscope (Apotome Zeiss AXIO Imager.Z1 with a 10x objective, ZEISS Plan-APOCHROMAT, NA: 0.45) to observe the injection sites, fiber, and virus expression.

To characterize the distribution of cortical neurons, the brains were cut at 50 µm intervals in sagittal sections (density profiles) or coronal sections (depth profiles) using a vibratome (Leica VT1200s). For density profiles, mosaic images (resolution of 1,023 µm/pixel) of slices with retrogradely labeled neurons were obtained using a fluorescence microscope (Zeiss AXIO Imagen.Z1) coupled to a digital camera (AxionCam MRm, 1.3 MP) with a 10x objective (ZEISS Plan-APOCHROMAT, NA 0.45). For depth profiles, mosaic images (0.66 µm/pixel) were acquired with a confocal microscope (Zeiss 780 LSM) using an LD PCI Plan-Apochromat 25x/0.8 Imm Corr DIC M27 objective.

Mosaic microscopy images were manually aligned in Amira software version 5.6.0 parallel to magnetic resonance imaging (MRI) images of the rat brain (Olivares-Moreno et al., 2017), using shared anatomical landmarks. Subsequently, references were placed at each marked neuron soma. To compute relative neuronal density profiles, distributions were obtained in steps of 250 µm x 250 µm for sagittal slices, while depth profiles were performed in steps of 50 µm along the vertical axis.

### Data analysis

#### Behavioral analysis

Shapiro-Wilk normality tests were used for the behavioral analysis during the training phases. For parametric data, a repeated measures ANOVA or paired t-test was applied. To evaluate learning, Bonferroni post-hoc test was used, considering the first training session as a control. The significance of differences in successes based on training session was determined (p < 0.05).

#### Lever trajectory

Lever trajectories were analyzed using codes specifically designed for the behavioral task in MATLAB Simulink® (version 2024a). Animals were categorized into three learning groups based on their lever trajectories: beginners (sessions 1-3), and experts (sessions 26-28). Subsequently, the following kinematic parameters for each category were evaluated (Supplementary Fig. 2A).

- Duration: total time to complete the lever movement.
- Amplitude: distance between the baseline and minimum lever point.
- Reaction Time: time between Go signal and lever press detection.
- Instantaneous velocity and acceleration.
- Push time: the time it takes to press the lever.

For kinematic parameter analysis, the Kolmogorov-Smirnov normality test was applied. For parametric data, a one-way ANOVA test followed by Bonferroni post-hoc test were computed. For non-parametric data, a Kruskal-Wallis test with Dunn’s post-hoc test or a Wilcoxon test were applied.

### Receiver operating characteristic (ROC) curve analysis

Lever trajectories during early or late phases of optogenetic photoinhibition were evaluated and compared with control lever trajectories. The lever trajectory averages and photoinhibition sessions were averaged for each animal to reconstruct the average trajectory. The comparisons were computed from all reconstructed individual trajectories. In this way, a time-by-time analysis was conducted, estimating the area under the ROC curve (AUROC) for each time interval (1.5 ms). Additionally, a permutation test randomly shuffling data at each interval for each trajectory was performed. AUROC values were considered statistically different if the real AUROC value was greater than 95% of the randomly shuffled values obtained (1000 permutations). This analysis identified the specific times when control trajectories differed from the trajectories recorded during photoinhibition.

## RESULTS

### Anatomical distribution of CS and CR neurons

To characterize the distribution of CR and CS neurons within the different layers of the cortex, FG retrograde tracer was injected into the red nucleus (AP: −5.7 mm, MD: 1 mm, DV: 6.8), and BDA was injected into the spinal cord (C4/C5 DV: 0.6 and 1.0 mm, ML:1 mm) of Wistar rats (250 to 300 g). We only analyzed experiments that had well-defined injection sites in the spinal cord and red nucleus (Figure 1 A-B). Coronal sections at AP: +1 mm with respect to bregma were analyzed. The vertical distribution of CS, CR, and double-labeled pyramidal neurons across the cortex (Fig. 1 C-D) was analyzed. The distribution of CS and CR neurons within layer 5 of the sensorimotor cortex (600 to 1200 µm in the motor cortex and 900 to 1300 µm in the somatosensory cortex) showed no separation between these two populations. This distribution remained consistent for the secondary motor cortex as well. On the other hand, the density of double labeled neurons was 10.6 ± 6.9 % in the motor cortex, 1.2 ± 0.4% in the secondary motor cortex, and 9.0 ± 6.5% in the somatosensory cortex. These are dual projection neurons, which project to the spinal cord and leave collaterals in the red nucleus.

**Figure 1.**
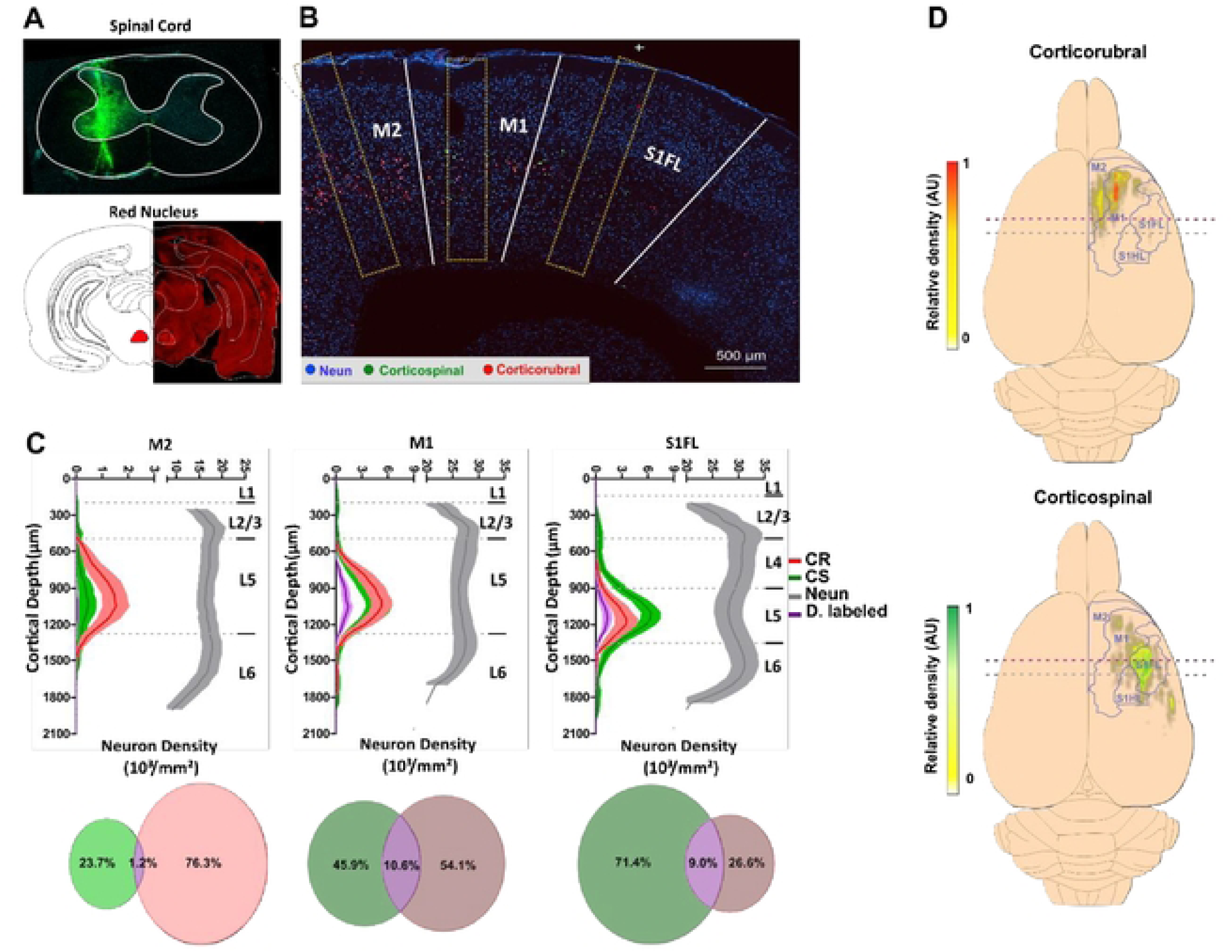
Anatomical distribution of CS and CR neurons in the cortex. Retrograde tracer injections (A); at the top, the site of spinal cord injection (BDA), and at the bottom, the site of injection in the red nucleus (FluoroGold). In (B), a coronal section of the cortex is shown (AP: +1 mm); in green, the soma of neurons labeled with BDA, with projections to the spinal cord; in red, the somas of neurons labeled with FluoroGold, with projections to the red nucleus; and in light blue, the somas of neurons labeled with Neun. Dotted boxes indicate the regions from which neuronal density profiles were obtained along the dorsoventral axis of the cortex (C); in green, the distribution of CS neurons; in red, the distribution of CR neurons; in gray, the distribution of neurons marked with Neun; and in purple, double-marked neurons. Shadows indicate standard error, and dashed vertical lines denote the different layers of the cortex. Venn diagrams show the percentage of CR and CS neurons in each cortex. In (D), density profiles show the relative density of CR neurons (above) and CS neurons (below). The gray dashed line at bregma. The magenta dashed line indicates the coordinate where depth distribution profiles were performed. SFL (anterior limb somatosensory cortex); M2 (secondary motor cortex); M1 (primary motor cortex); SHL (posterior limb somatosensory cortex).

There is a difference in the anteroposterior and mediolateral distribution of the two populations of interest. CS neurons exhibited a higher density in somatosensory regions (S1FL 71.4 %, while CR neurons showed a greater density in motor regions (M1 54.1 % and M2 76.3 %) (Fig. 1 C: Venn diagrams, Fig. 1 D, and Supplementary Fig. 1).

### Behavior

In order to test the role of CS and CR neurons during motor execution, rats were trained to press a lever. The operant conditioning task was standardized in freely moving Wistar rats (n=20) under water restriction (water access two times per day: during training and one-hour post-training)). The animals learned to press a lever after a cue light signal was turned off (Fig. 2 A). The learning curve shows that animals reached a 55 % success rate around session 26, but a significant difference was observed from session 6 (repeated measures ANOVA: F=20.88, p<0.0001; Bonferroni session 1 vs session 6: t=4.861, p≤0.0001) (Fig. 2 B). The observed hits demonstrated a significant increase in the performance of beginner (sessions 1 to 3) and expert (sessions 26 to 28) rats, indicating that the task had been learned (Fig. 2 B and D; n = 20; Student’s t-test: t = 13.82, df = 94; p = 0.0001).

**Figure 2.**
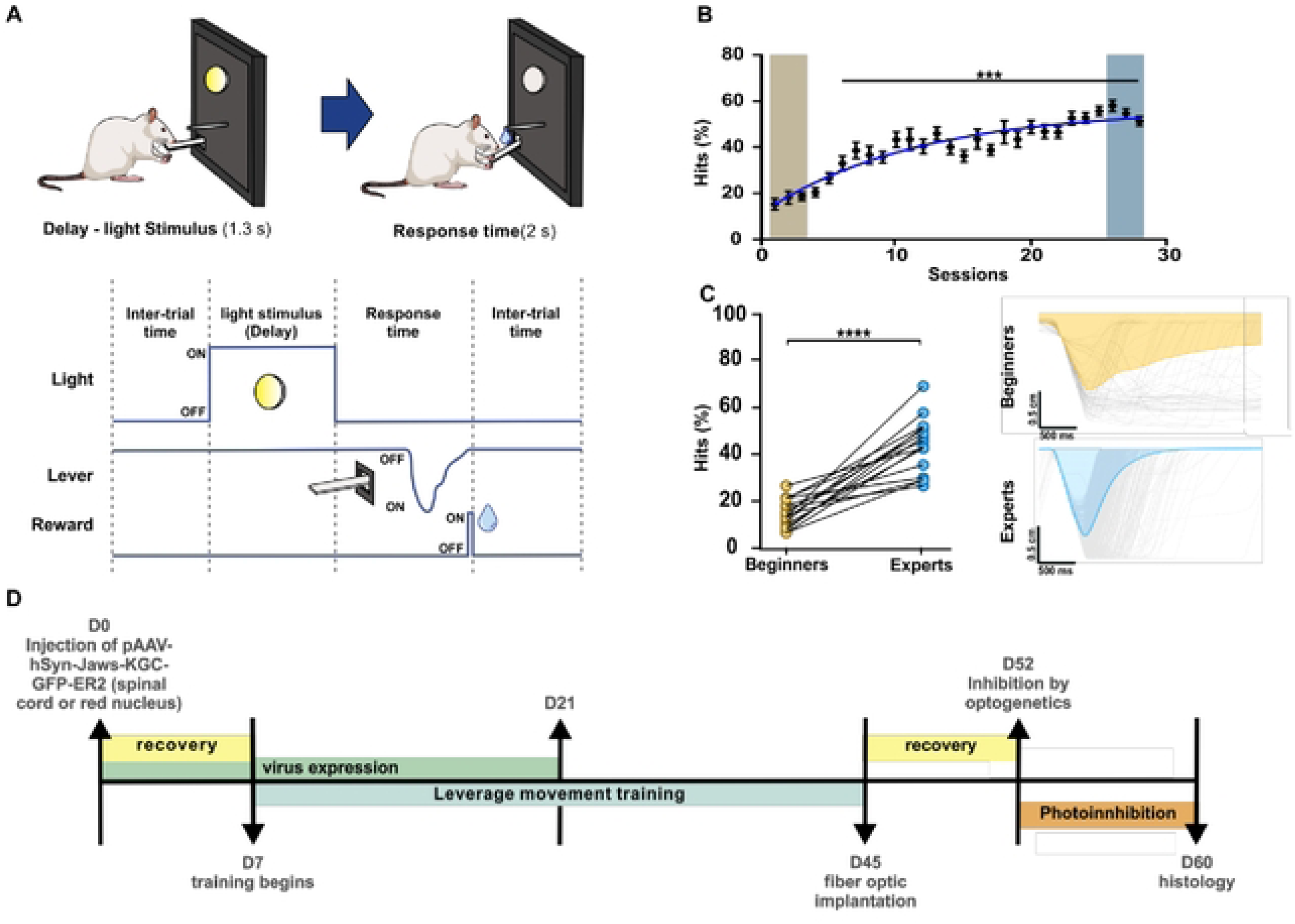
Behavioral paradigm. In (A), light stimulus or delay (700 - 1300 ms), response time (2000 ms), and inter-trial interval (2000 - 4000 ms) are illustrated. Learning curve, n=20 (B), with vertical lines indicating standard error. In (C), comparison of hits between beginner (sessions 1-3) and expert rats (sessions 26-28) (n = 20), showing lever presses for one session per condition on the right, individual trajectories in gray lines, and the average trajectory in shaded line. In (D), experimental design timeline.

We found an improvement in expert animals, as their lever presses became more stereotyped in (Fig 2 C, right), with a notable decrease in duration (Supplementary Fig 2 H; n=16; Wilcoxon W= 937000; p < 0.0001; mean beginners: 0.91 ± 0.27 s; mean experts: 0.73 ± 0.29 s), push time (Supplementary Fig 2 G; n=16; Wilcoxon W= 591200; p < 0.0001; mean beginners: 0.40 ± 0.14 s; mean experts: 0.31 ± 0.13 s), reaction time (Supplementary Fig 2 J; n=16; Wilcoxon W= 487900; p < 0.0001; mean beginners: 0.822 ± 0.617 s; mean experts: 0.553 s ± 0.470) and amplitude (Supplementary Fig 2 H; n=16; Wilcoxon W= 866200; p < 0.0001; mean beginners: 1.35 ± 0.18 cm; mean experts: 1.28 ± 0.21 cm). We also observed that parameters such as push speed (Supplementary Fig 2 C; n=16; Wilcoxon W= 1088000; p < 0.0001; mean beginners: −1.11 ± 0.94 cm/s; mean experts: −1.92 ± 1.16 cm/s) and pull speed (Supplementary Fig 2 D; n=16; Wilcoxon W= −213000; p < 0.0001; mean beginners: 1.34 ± 1.17 cm/s; mean experts: −2.27 ± 1.39 cm/s) increased with practice.

### Optogenetic inhibition of CS and CR neurons

After training, optogenetic inhibition protocols were conducted by stimulating rats previously injected with pAAV-hSyn-Jaws-KGC-GFP-ER2 in the spinal cord or red nucleus (Fig. 2D; n=8). As previously observed with retrograde tracers, viral expression in CS neurons is more predominant in the motor cortex at AP + 2.20 (Fig. 3 B and C, left). The CR neurons (Fig. 3A, right) are expressed in the motor cortex but with a higher density in the secondary motor cortex at AP + 2.20 (Fig. 3 B and C). Thus, the optical fiber was implanted in the motor cortex region after training (Fig. 2 D and Supplementary Fig. 3). Three kinds of trials were conducted in a training session: first, a control trial with no photoinhibition; second, an early inhibition trial, where optogenetic inhibition was applied during the light stimulation period; and third, a late inhibition trial, where optogenetic inhibition was applied during the animal’s response period (Fig. 3 D, below).

**Figure 3.**
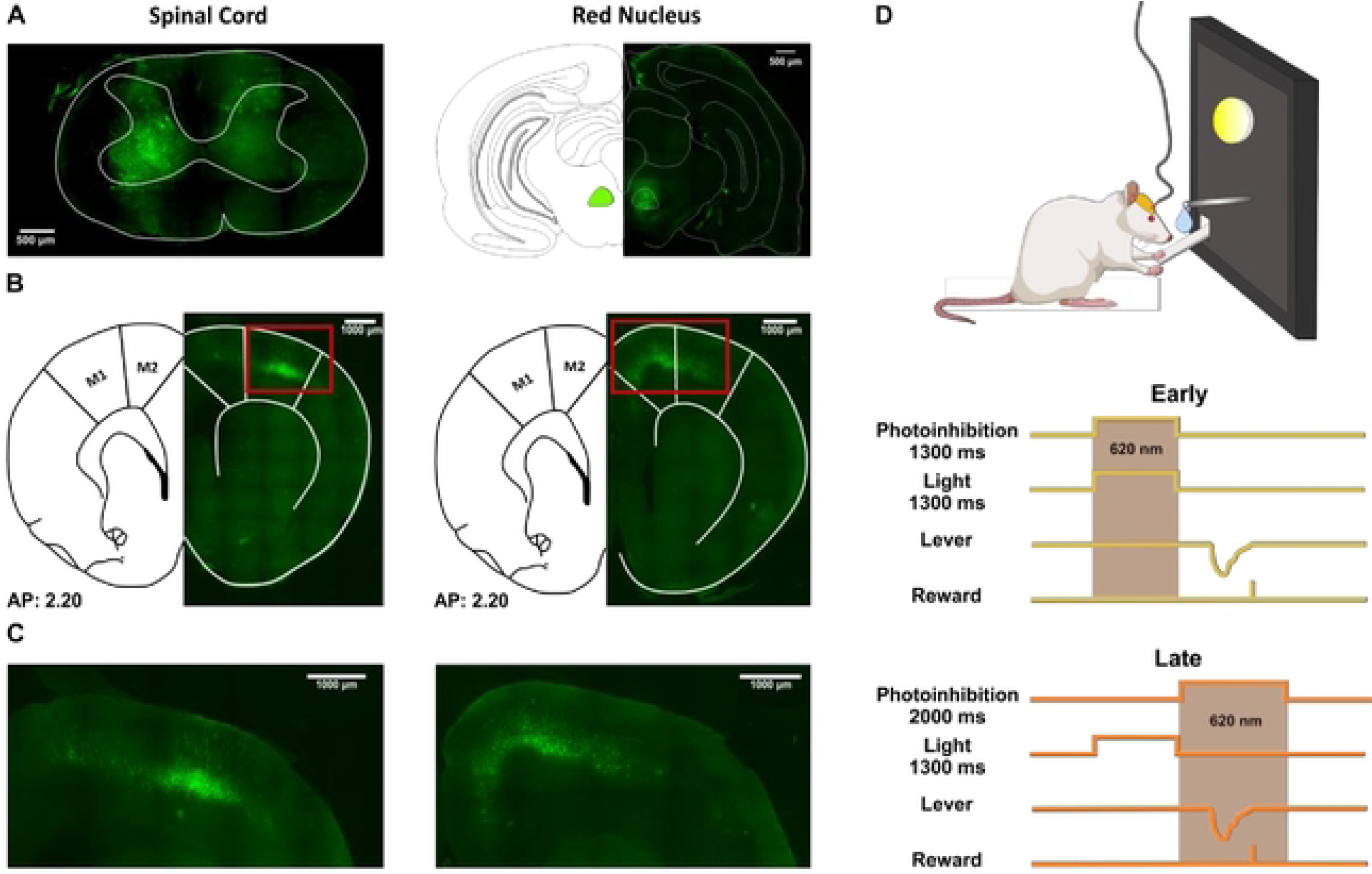
Expression of the virus pAAV-hsyn-Jaws-KGC-GFP-ER2 and photoinhibition protocol. In (A), injection sites in the spinal cord at C5 level (left) and in the red nucleus (right). In (B), virus expression in cortices (Bregma 2.20) M1 and M2 for CS and CR neurons. In (C), amplification of the expression site (red box in B) of CS and CR neurons. In (D), inhibition protocol during early phase and late phase; shaded region indicates the time of inhibition.

No significant differences were found between the hits performed by CR neuron groups for either of the inhibition protocols (Fig. 5 B; CR: n = 4; repeated measures ANOVA F = 2.357; p = 0.1182; mean control: 46.86 ± 35.13 %; mean early: 28.71 ± 11.02 %; mean late: 31.69 ± 12.30 %). However, significant differences were found between the hits in the CS neuron group for early inhibition (Fig. 5 B; CS: n = 4; repeated measures ANOVA F = 9.306; p = 0.0004; Bonferroni control vs early p<0.05; mean control: 45.66 ± 13.85 %; mean early: 24.21 ± 16.03 %; mean late: 33.91± 15.62 %).

On the other hand, the movement kinematics were altered in the rats with inhibition of M1 CS neurons. Early inhibition resulted in increased lever pressing duration (Fig. 4C; n = 4; Kruskal-Wallis statistic = 23.53; p < 0.0001; Dunn control vs early p < 0.05; mean control: 0.81 s ± 0.33; mean early: 0.94 s ± 0.36; mean late: 0.87 ± 0.37 s). However, neither the amplitude nor the reaction time showed significant changes in the two inhibition protocols compared to the control for this group of rats (Fig. 4B; n = 4; Kruskal-Wallis statistic = 5.667; p= 0.0588; mean control: 1.23 cm ± 0.25; mean early: 1.27 ± 0.26 cm; mean late: 1.25 ± 0.24 cm; Fig. 4 D; Kruskal-Wallis statistic = 7.317; p= 0.0258; mean control: 0.637 ± 0.508 s; mean early: 0.740 ± 0.600 s; mean late: 0.548 ± 0.548 s).

**Figure 4.**
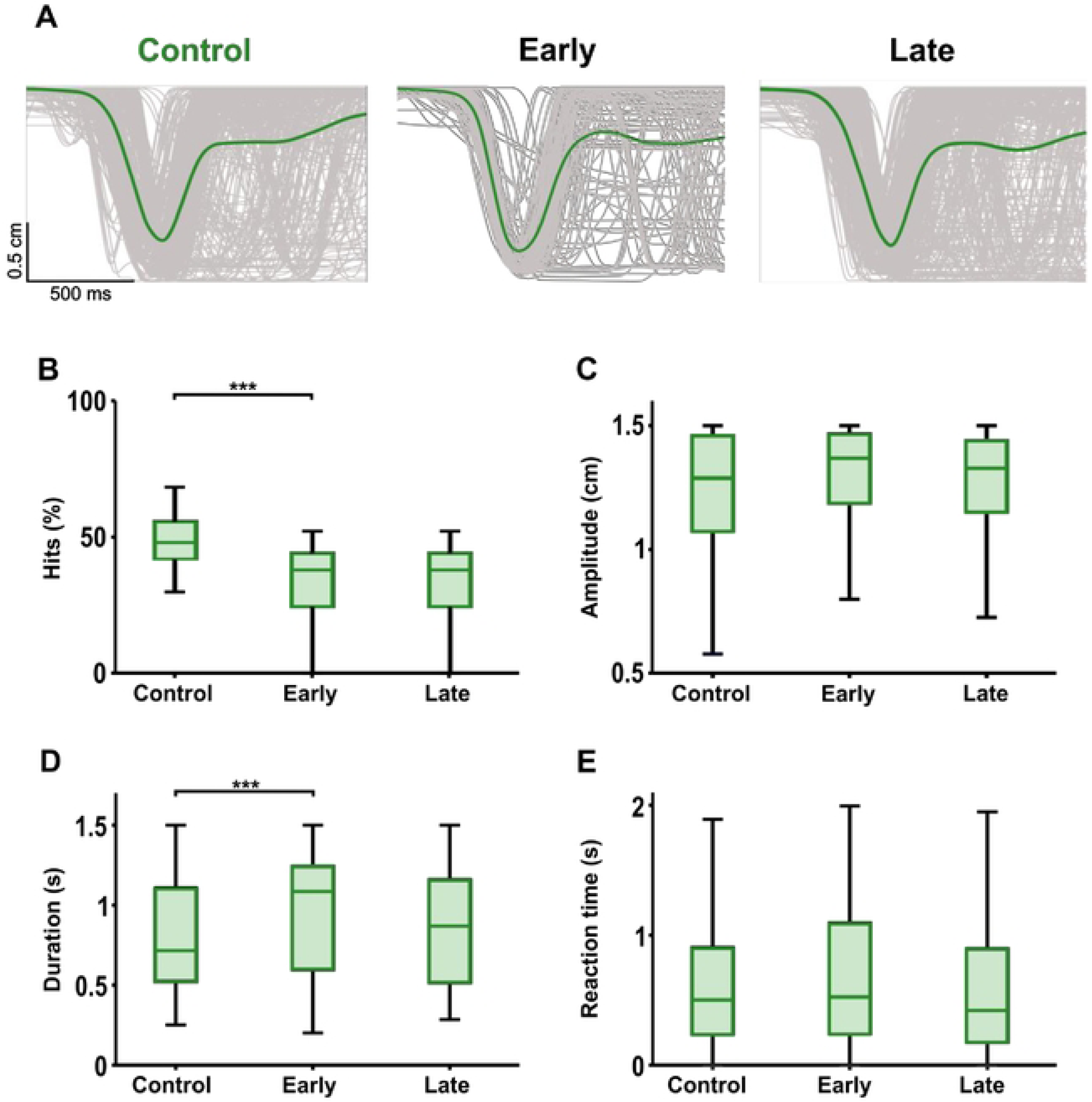
Effect of CS neuron inhibition on lever pressing trajectories. (A), lever trajectories of the CS neuron group upon inhibition. Averaged control lever press trajectories are shown in the solid green line under control conditions (left), during early inhibition (middle), and during late inhibition (right) is presented. Individual lever presses are shown in gray, while the average of lever presses is shown in color. Different kinematic parameters of lever pressing movement were evaluated: hits (B), amplitude (C), duration (D), reaction time (E). Comparisons were made between control lever presses vs early inhibition and control vs late inhibition (n =4; Kruskal Wallis with Dunn’s post hoc test).

Push speed, push acceleration, and push time were not significantly altered in either type of inhibition for CS neurons compared to their controls (Supplementary Fig. 4A; n = 4; Kruskal-Wallis test = 5.107; p= 0.0778; mean control: −1.58 cm/s ± 1.04; mean early: −1.52 ± 1.11 cm/s; mean late: −1.66 ± 1.00 cm/s; Supplementary Fig. 4 C; Kruskal-Wallis test = 14.76; p= 0.0006; mean control: 1.51×10^-2^ 2 ± 2.06×10^-2^ cm/s; mean early: 1.30 ×10^-2^ 2 ± 2.25 ×10^-2^ cm/s; mean late: 1.85 ×10^-2^ 2 ± 2.09 ×10^-2^ cm/s; Supplementary Fig. 4E; n = 4; Kruskal-Wallis test = 0.0361; p= 6.641; mean control: 0.31 ± 0.15 s; mean early: 0.36 ± 0.19 s; mean late: 0.34 ± 0.19 s). Nevertheless, pull velocity and pull acceleration showed significant differences in both early and late inhibition with respect to their controls in the group of CS neurons (Supplementary Fig. 4B; n = 4; Kruskal-Wallis statistic = 17.28; p= 0.0002; Dunn control vs early p < 0.05; Dunn test control vs late p < 0.05; mean control: 2.07 ± 1.32 cm/s; mean early: 2.54 ± 1.36 cm/s; mean late: 2.42 ± 1.29 cm/s; Supplementary Fig. 4D; n=4; Kruskal-Wallis test = 17.36; p= 0.0002; Dunn control vs early p < 0.05; Dunn control vs late p < 0.05; mean control: 1.90 x 10^-2^ ± 2.72 x 10^-2^ cm/s^2^; mean early: 2.75 ×10^-2^ ± 2.81 x 10^-2^ cm/s^2^; mean late: 2.80 ×10^-2^ ± 2.35 x 10^-2^ cm/s^2^).

On the other hand, CR neuron group showed alterations in kinematic parameters mostly during early inhibition (Fig. 5). In contrast, the amplitude of the lever pressing decreased during late and early inhibition (Fig 5B; n = 4; Kruskal-Wallis statistic = 9.054; p < 0.0108; Dunn control vs early p < 0.05; Dunn control vs late p < 0.05; mean control: 1.32 ± 0.22 cm; mean early: 1.26 ± 0.27 s; mean late: 1.28 ± 0.25 s). Conversely, the duration and reaction time increased during the early inhibition of CR neurons (Fig 5C; n = 4; Kruskal-Wallis statistic = 28.21; p < 0.0001; Dunn control vs early p < 0.05; Dunn control vs late p < 0.05; mean control: 0.69 ± 0.26 s; mean early: 0.73 ± 0.25 s; mean late: 0.64 ± 0.22 s; Fig. 5 D; Kruskal-Wallis statistic = 19.54; p < 0.0001; Dunn control vs early p < 0.05; mean control: 0.600 ± 0.499 s; mean early: 0.746 ± 0.542 s; mean late: 0.656 ± 0.524 s).

**Figure 5.**
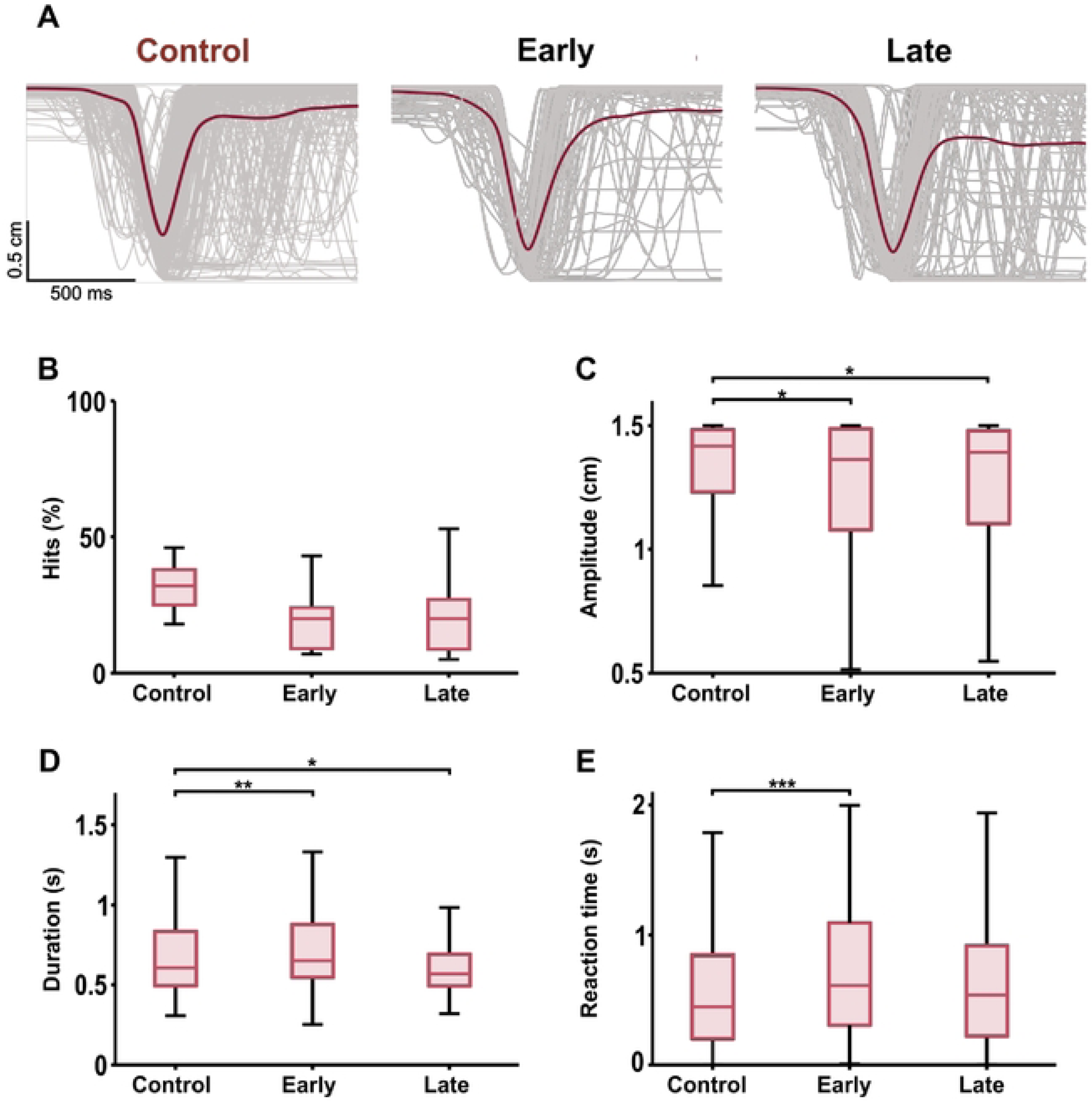
Effect of CR neuron inhibition on lever pressing trajectories. (A), lever trajectories of the CR neuron group upon inhibition. Averaged control lever press trajectories are shown in the solid red line under control conditions (left), during early inhibition (middle), and during late inhibition (right). Individual lever presses are shown in gray, while the average of lever presses is shown in color. Different kinematic parameters of lever pressing movement were evaluated: hits (B), amplitude (C), duration (D), reaction time (E). Comparisons were made between control lever presses vs early inhibition and control vs late inhibition (n =4; Kruskal Wallis with Dunn’s post hoc test).

When evaluating the effect of inhibiting CR neurons in the motor cortex, we observed a significant decrease in push and pull speeds compared to the control (Supplementary Fig. 5A; n = 4; Kruskal-Wallis = 73.74; p< 0.0001; Dunn control vs early p < 0.05; Dunn’s control vs late p < 0.05; mean control: −1.52 ± 0.77 cm/s; mean early: −1.20 ± 0.75 cm/s; mean late: −1.19 ± 0.58 cm/s; Supplementary Fig. 4B; n=4; Kruskal-Wallis statistic = 58.64; p< 0.0001; Dunn control vs early p < 0.05; Dunn control vs late p < 0.05; mean control: 1.73 ± 0.93 cm/s; mean early: 1.34 ± 0.88 cm/s; mean late: 1.35 ± 0.90 cm/s), as well as a decrease in pull acceleration (Fig. 4D; n=4; Kruskal-Wallis test = 46.70; p< 0.0001; Dunn control vs early p < 0.05; Dunn control vs late p < 0.05; mean control: 1.40×10^-2^ ± 2.28 x 10^-2^ cm/s^2^; mean early: 0.58 x 10^-2^ ± 2.01 x 10^-2^ cm/s^2^; mean late: 0.46 x 10^-2^ ± 1.77 x 10^-2^ cm/s^2^). These effects were seen with both types of inhibition. However no significant changes were observed in push acceleration (Fig. 4 D; n=4; Kruskal-Wallis statistic = 14.76; p= 0.0006; mean control: 0.96 x 10^-2^ ± 2.11 x 10^-2^ cm/s^2^; mean early: 0.66 ×10^-2^ ± 1.42 x 10^-2^ cm/s^2^; mean late: 0.70 ×10^-2^ ± 1.39 x 10^-2^ cm/s^2^). Finally, push time increased significantly during early inhibition in CR M1 neuron group (Fig 5D; n=4; Kruskal-Wallis statistic = 31.40; p < 0.0001; Dunn control vs early p < 0.05; mean control: 0.28 ± 0.09 s; mean early: 0.32 ± 0.11 s; mean late: 0.29 ± 0.08 s). No significant changes in the kinematics parameters were observed when a 465 nm light was applied to CS or CR neurons instead of the 620 nm light.

### Temporal trajectory analysis

A bin-by-bin ROC curve analysis was conducted to determine which epochs of the task showed differences between the individual control trajectories and the individual trajectories of the trials with inhibition. Trajectories of animals from the CS (Fig. 6A and Fig. 4A) and CR neuron groups (Fig. 6B and Fig. 5A) exhibit significant differences during early and late inhibition. In CS neurons, these differences were more evident after movement onset and persisted until the end (flexion phase) (Fig. 6A, upper part; ROC curve analysis p < 0.05). When CR neurons were inhibited, the differences were observed mainly in the pre-movement epoch and persisted throughout all movements (Fig. 6B, lower part; ROC curve analysis p < 0.05).

**Figure 6.**
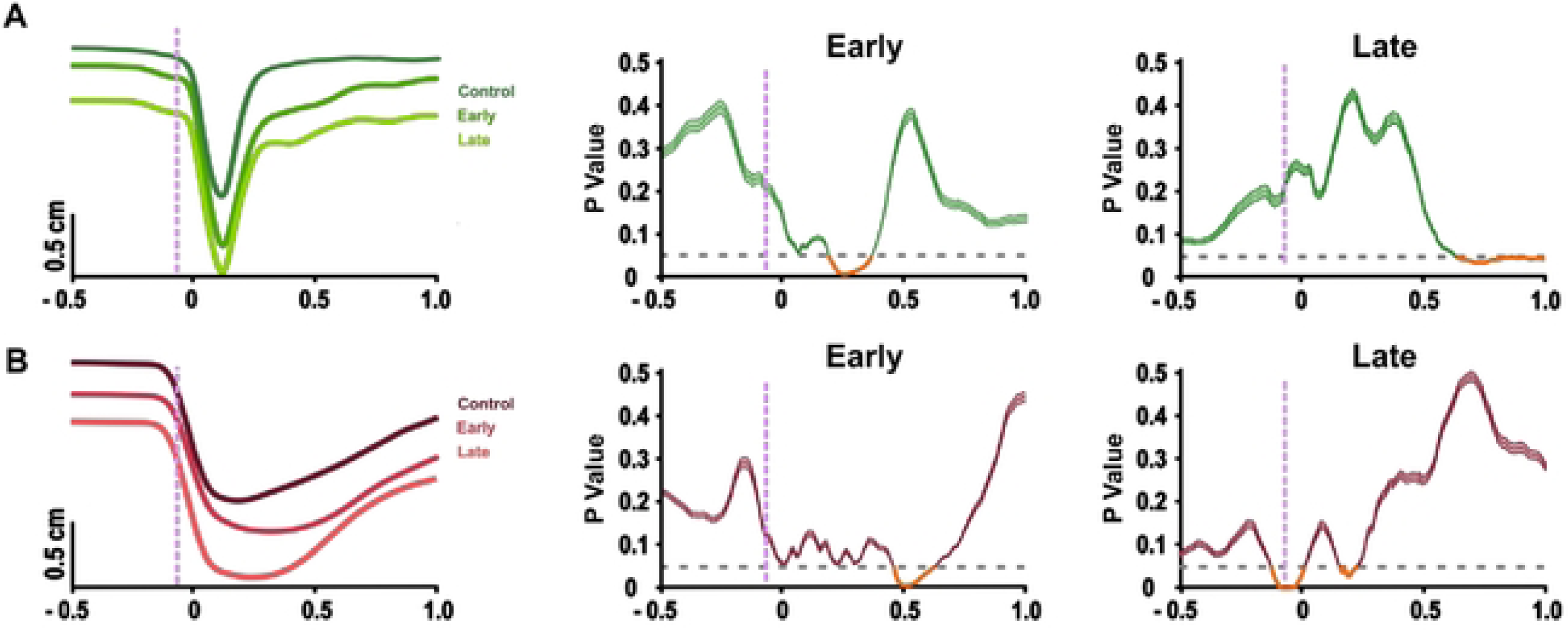
Kinematic parameters of lever pressing trajectory between inhibited and control trials. (A), averaged lever press trajectories of CS inhibited animals (Left graphs). The solid green line under control conditions (upper), during early inhibition (middle), and during late inhibition (lower). The averaged p-value (±SE) for all rats with early (middle graph) and late (right graph) inhibitions in CS neurons. Significant values, highlighted in orange, were identified in the ROC curve analysis throughout the temporal window (p-value < 0.05). The gray horizontal dotted line indicates the threshold of 0.05, while the magenta vertical line marks the beginning of the leverage movement. (B), the same as A but for CR inhibited rats.

## DISCUSSION

In this study, we used an operant conditioning model in which rats had to press a lever guided by a light stimulus. The protocol allowed us to assess movement preparation (light stimulus) and execution (Go signal). The learning curve shows that animals improved their performance from the 6th training session, achieving a success rate of 55 % (Fig. 2B). A significant difference in success rates between beginner and expert rats was observed from the first to the last training session, indicating that the animals progressed through the perceptual and associative learning phases. Additionally, an analysis of kinematic parameters revealed decreases in duration, reaction time, and movement amplitude, suggesting that movements became more precise and stereotyped in later training stages (Laubach et al., 2000; Makino et al., 2016; Papale & Hooks, 2018). This enabled us to assess the involvement of CS and CR pathways at different stages of the movement.

The role of the cortex in motor control is well documented. As a more evolutive recent structure compared to subcortical motor nuclei (Olivares-Moreno et al., 2021), the cortex modulates the spinal cord directly via the CS pathway and indirectly via projections to older motor nuclei like the red nucleus (CR pathway). These pathways are thought to be involved in different stages of movement, including planning and execution (Houk, 1991; Isomura et al., 2009; Li et al., 2015).

### Anatomical distribution

The functional diversity of neurons in layer 5 of the cortex, which are the primary output information from the cortex, is still debated. One view suggests that these neurons, which project to multiple subcortical structures (Guo et al., 2017; Kita & Kita, 2012), play a role in associative movement control without clear compartmentalization of motor commands. This implies low functional diversity, with the same information sent to various movement-related structures simultaneously. Conversely, another perspective suggests that layer 5 neurons exhibit high functional diversity based on their specific projection targets (Economo et al., 2018; Isomura et al., 2009; Narayanan et al., 2017; Rojas-Piloni et al., 2017). This last viewpoint supports the idea that the cortex regulates older motor systems and pathways, allowing for distinct control of movements by different neuronal subgroups (Olivares-Moreno et al., 2021).

Our histological findings are consistent with the latter proposal, as they show an topographic segregation of the two evaluated populations (CR and CS). CR neurons are more densely located in motor regions like M1 and M2, while CS neurons are more concentrated in S1 and M1, with less presence in S2 and no presence in M2 (Fig. 1 D). This distribution aligns with findings from other studies, which show that there is a differential distribution of layer 5 projection neurons depending on their target site (Akintunde & Buxton, 1992; Economo et al., 2018; Lopez-Virgen et al., 2022; Olivares-Moreno et al., 2017). Studies, such as those by Kita and Kita (2012), have shown that the subthalamic nucleus of rats mainly receives long-range axon collaterals with multiple subcortical targets. Similarly, Guo found that layer V type I neurons in mice have a high probability of projecting to multiple regions, with the thalamus and midbrain being the most frequent targets (Guo et al., 2017; Kita & Kita, 2012). Unlike previous research that reported cortical projection neurons sending collaterals to multiple subcortical structures, this study found a low percentage (about 10%) of double-labeled neurons. Our data are similar to those reported by Akintunde (Akintunde & Buxton, 1992), who found approximately 4% of neurons doubly labeled with CR and CS. The discrepancies between the studies may be related to the methodologies used for the measurements, which in some cases involved viral infections and in others retrograde markers (Akintunde & Buxton, 1992; Guo et al., 2017; Kita & Kita, 2012).

### Inhibition of CS and CR M1 neurons

For this study, we injected a retrograde virus into the spinal cord and red nucleus to analyze the role of these pathways during movement (Fig. 3 A). In addition, we applied optogenetic inhibition protocols to investigate their effects on movement phases (Fig. 3 D). Fibers were implanted in the M1 where these neuron populations converge (Supplementary Fig. 3).

No significant change in the success rate was observed in any of the types of inhibition in the CR group of animals compared to the control (Fig. 5B). This suggests that the animals performed the task successfully despite the inhibition. Other studies have reported more pronounced effects with lesions or total inhibitions (Ishida et al., 2016, 2019). However, differences associated with movement preparation were found in the CS group during early inhibition. This finding is consistent with the function of CS neurons projecting to the dorsal horn, which may be involved in modulating sensory and proprioceptive inputs (Macías et al., 2022; Olivares-Moreno et al., 2017) (Fig. 4B). Here, the ability to inhibit CS and CR neurons is restricted by the numerical aperture of the fiber used, which limits the effect to a radius of 200 micrometers. Therefore, it is not possible to observe an effect as drastic as that reported in studies where complete lesions or inhibitions of the pathways were performed (Ishida et al., 2016, 2019). Additionally, although the lever movement was still performed, changes in execution were observed, prompting an analysis of various kinematic parameters to identify these alterations.

Early and late inhibition of CS neurons increased the duration of lever pressing (Fig. 4C), pull velocity (Supplementary Fig. 4 B), and pull acceleration (Supplementary Fig. 4D) compared to the controls. The changes observed when inhibiting a preparatory phase and an execution phase can be explained by the fact that the CS tract has distinct neuronal subgroups that modulate sensory information and motor execution (Lemon, 2008; Olivares-Moreno et al., 2017), and different subtypes of spinal cord interneurons (Lemon, 2008). Thus, the CS tract could play a key role in sensorimotor integration by modulating the synaptic noise into the spinal cord and receiving motor commands (Moreno-López et al., 2016; Olivares-Moreno et al., 2017; 2019; Soteropoulos, 2018).

These changes in CS inhibition are consistent with the observed alterations in animal trajectories (Fig. 6A), suggesting that the longer lever press duration and increased release speed and acceleration may indicate a decrease in control over the lever return movement (flexion movement). In this way, Fetz et al. have shown that the CS tract facilitates both flexor muscles (51 %) and extensor muscles (48 %) (Fetz, 1993; Massion, 1967). However, the minimal effect on lever execution observed with CS neuron inhibition aligns with findings that CS neuron involvement in movements changes with training (Macías et al., 2022). In expert animals, CS neurons in area M1 are active before and during movement execution, but their role may decrease in highly trained movements (Peters et al., 2017). This indicates that the cortex becomes less involved in executing well-learned movements and only re-engages to make corrections in response to external disturbances when needed (Hwang et al., 2021; Mathis et al., 2017; Wolff et al., 2022).

Early and late inhibition of CR neurons significantly affected lever execution. Early inhibition increased the duration of lever pressing (Fig. 5C) while significantly decreasing amplitude (Fig. 5B), push and pull speeds, and push and pull accelerations. These effects are in line with changes observed in trajectory performance (Fig. 6B), indicating issues with initiating and executing the movement, including extension (lever press) and flexion (lever release) phases. Some studies have linked damage to the CR tract and disinhibition of the red nucleus with forelimb extension problems (Basile et al., 2020). However, these studies have not been replicated. On the other hand, it has been demonstrated that the rubrospinal tract controls both extensor and flexor muscles (Fetz, 1993; Massion, 1967), which is consistent with the results observed in our study.

Early phase inhibition may be linked to CR tract modulation of red nucleus activity, which indirectly affects the rubrospinal tract. This modulation probably regulates the excitability of signals from the contralateral interposed nucleus of the cerebellum (Canedo, 1997; Massion, 1988). Thus, the CR tract not only aids in the initiation and termination of voluntary movements but also modulates the red nucleus’s basal activity (Canedo, 1997; Massion, 1988), which is crucial for proper movement execution. Additionally, early-phase CR neuron inhibition increased reaction times (Fig. 5D), which is consistent with findings in cats where red nucleus neurons are active during reaction times before lever release (Amalric et al., 1983).

Inhibition of CR neurons during the late phase caused a significant decrease in parameters such as movement duration (Fig. 5 C), amplitude (Fig. 5 B), push and pull speed (Supplementary Fig. 5 A-B), pull acceleration (Supplementary Fig. 5 D), and descent time (Supplementary Fig. 5 E). Similar results have been observed when inhibiting M2 and M1 cortices in mice, disrupting the proper execution of a reaching movement (Galiñanes et al., 2018). On the other hand, training has been found to strengthen the association between M2 and M1 areas, as well as their relationship to motor performance (Veuthey et al., 2020). Considering the high population density of CR neurons in these two areas (Fig. 1 D), it could be argued that this neuronal population is involved in both the preparation and execution of movements.

### Functional differences

The convergence of CS and rubrospinal projections on their targets within the spinal cord, modulating local interneurons and propriospinal neurons (Alstermark et al., 1981; Illert et al., 1976, 1977), has led to the suggestion that there is a close relationship between these two motor pathways. Additionally, it has been documented that when there is damage to the red nucleus or loss of the cortico-rubrospinal pathway, movement execution is impaired, although there may be some compensation from the CS tract. Similarly, plasticity has been observed in the cortico-rubrospinal tract when there is damage to the CS tract (Humphrey & Rietz, 1976; Ishida et al., 2016, 2019; Satoh et al., 2007; Williams et al., 2014). In both cases, recovery is incomplete, suggesting that they are complementary pathways for transmitting information rather than copies of the same motor command.

Considering that the cortex and the parvocellular region of the red nucleus (CR target) emerged almost simultaneously during evolution, perhaps in response to the need for better limb control and increased complexity of movements beyond locomotion and “gross” movements, it raises the question of how these new structures and the preexisting ones are reorganized for motor control (Olivares-Moreno et al., 2021). Kennedy (1990) proposes that the CS system, is predominantly involved in the learning of new movements, nonetheless, the cortico-rubro-olivary tract and the rubrospinal tract more with the proper execution of learned or automated movements. Yet, the animals we evaluated are experts, which explains the differences in the involvement of the two pathways, with more evident changes during CR neuron inhibition (Kennedy, 1990; Kennedy & Humphrey, 1987; Kuypers & Lawrence, 1967).

Our study demonstrated that CR and CS neurons, subpopulations of PTNs, play an essential role in motor performance by modulating various kinematic parameters. These findings suggest that these neuronal populations contribute differently to sensorimotor integration, indicating that the cerebral cortex can reorganize neural circuits to execute a previously learned movement, even under inhibition conditions (Mathis et al., 2017). In this regard, it is suggested that these two pathways are necessary for proper preparation and execution of a movement.

## ABBREVIATIONS

PTNs: Pyramidal tract neurons;
CR: corticorubral;
CS: corticospinal.

## ACKNOWLEDGEMENTS

RM-P and LV_J are doctoral students from the Programa de Doctorado en Ciencias Biomédicas, Universidad Nacional Autónoma de México (UNAM) and received fellowships from CONAHCyT (934973 and 1094565). This work was supported by grant PAPIIT-DGAPA IN201624. We thank Jessica Gonzalez Norris for proofreading the manuscript and Cutberto Dorado, Nydia Hernandez, Ericka de los Rios, Alejandra Castilla, and Martín Garcia Servin, Christian Josué Delgado Guzmán, Edgar Bolaños Aquino for providing technical assistance.

## FIGURE LEGENDS

**Supplementary Figure 1.**
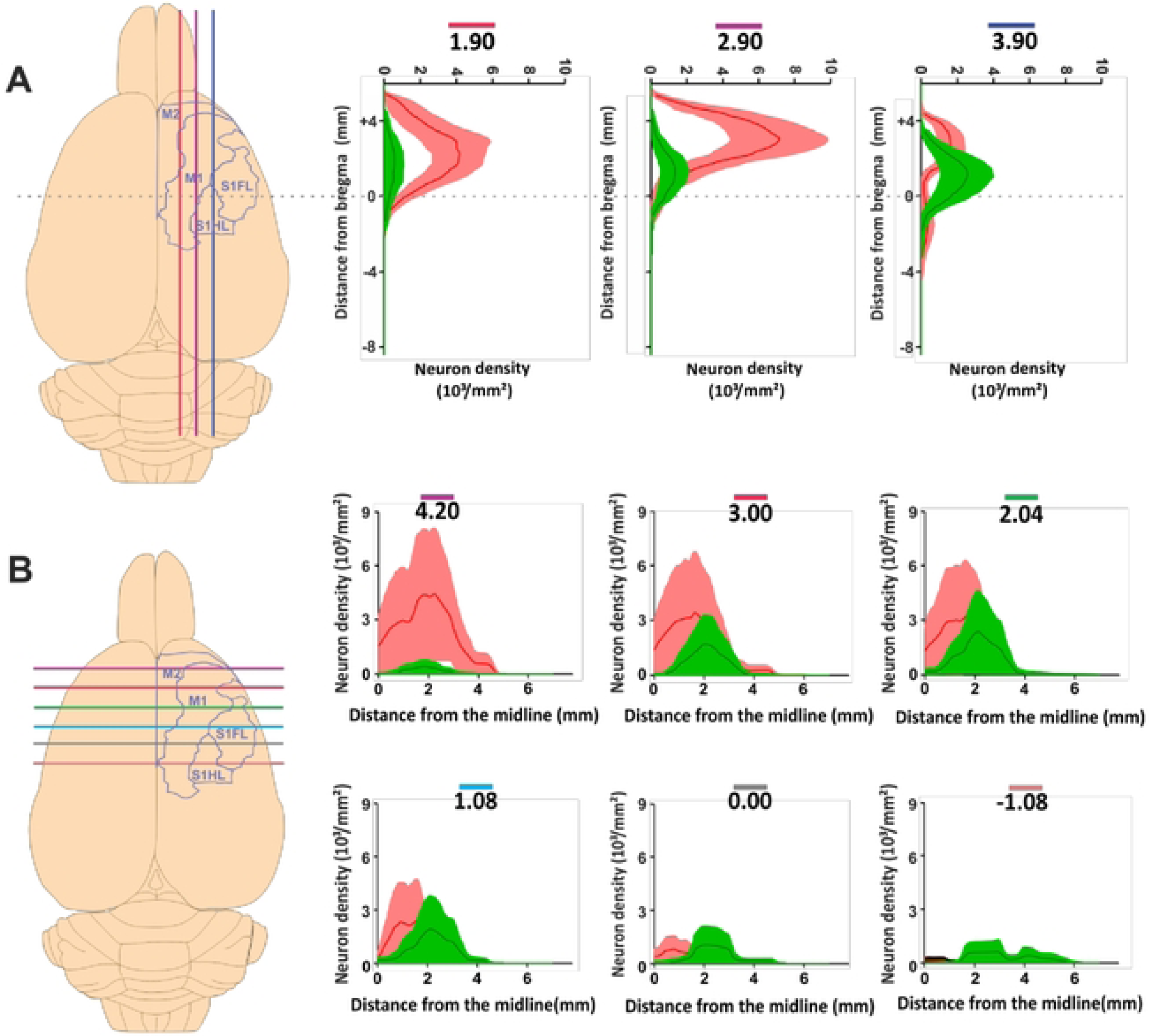
Anteroposterior and mediolateral distribution of CS and CR neurons in the cortex. In (A), mediolateral distribution of CS and CR neurons. In (B), anteroposterior distribution of the CS and CR neurons. In red, the distribution of CR neurons; in green, the distribution of CS neurons. Shadows indicate standard error. The colored bars indicate the mediolateral and anteroposterior coordinates where the analyses were performed.

**Supplementary Figure 2.**
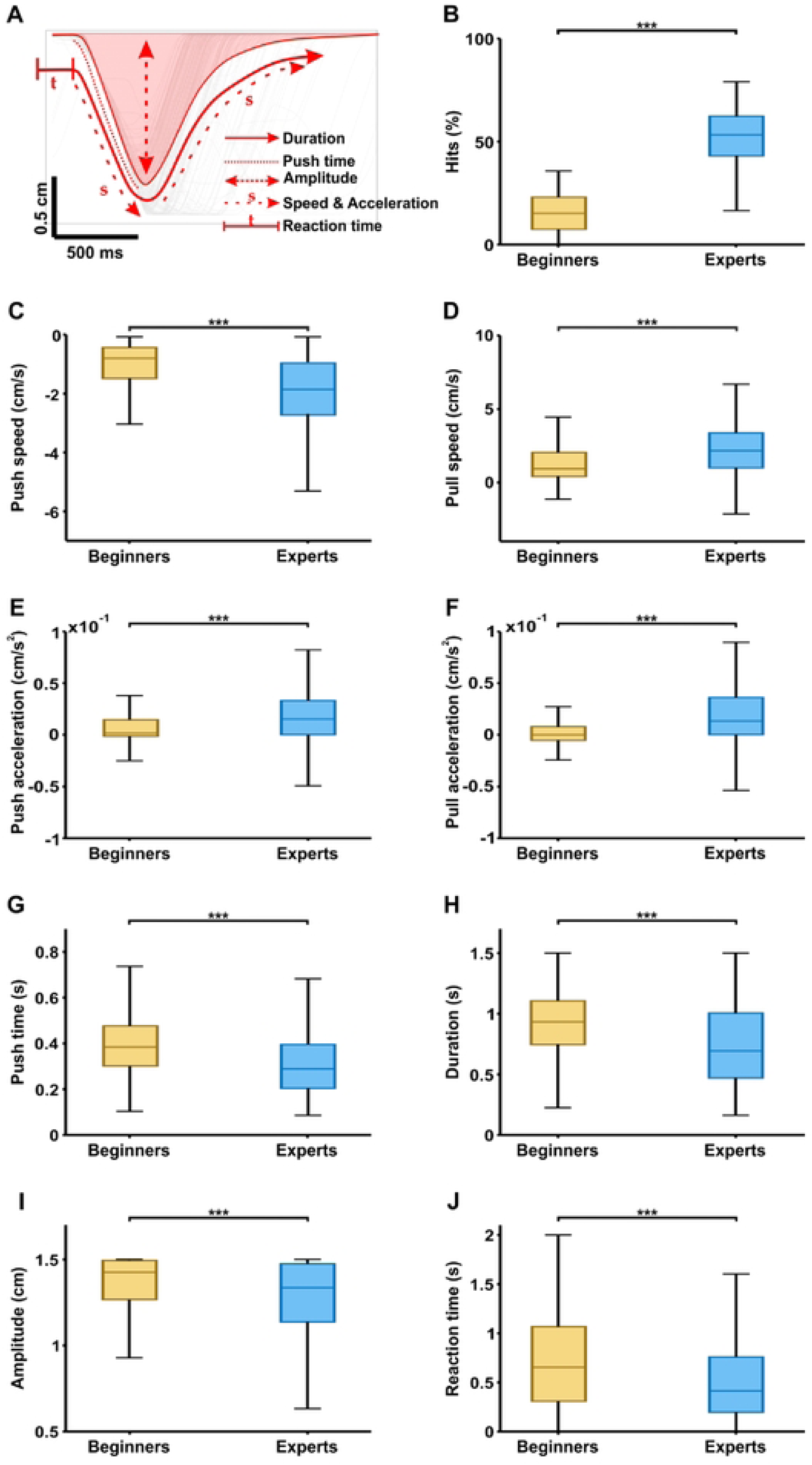
Kinematic parameters of lever pressing trajectory during learning. (A) depicts a graph of evaluated kinematic parameters. Hits (B), push speed (C), pull speed (D), push acceleration (E), pull acceleration (F), push time (G), duration. Movement (H), amplitude (I), reaction time (J). (n = 20; Wilcoxon. *** <0.001, *<0.05).

**Supplementary Figure 3.**
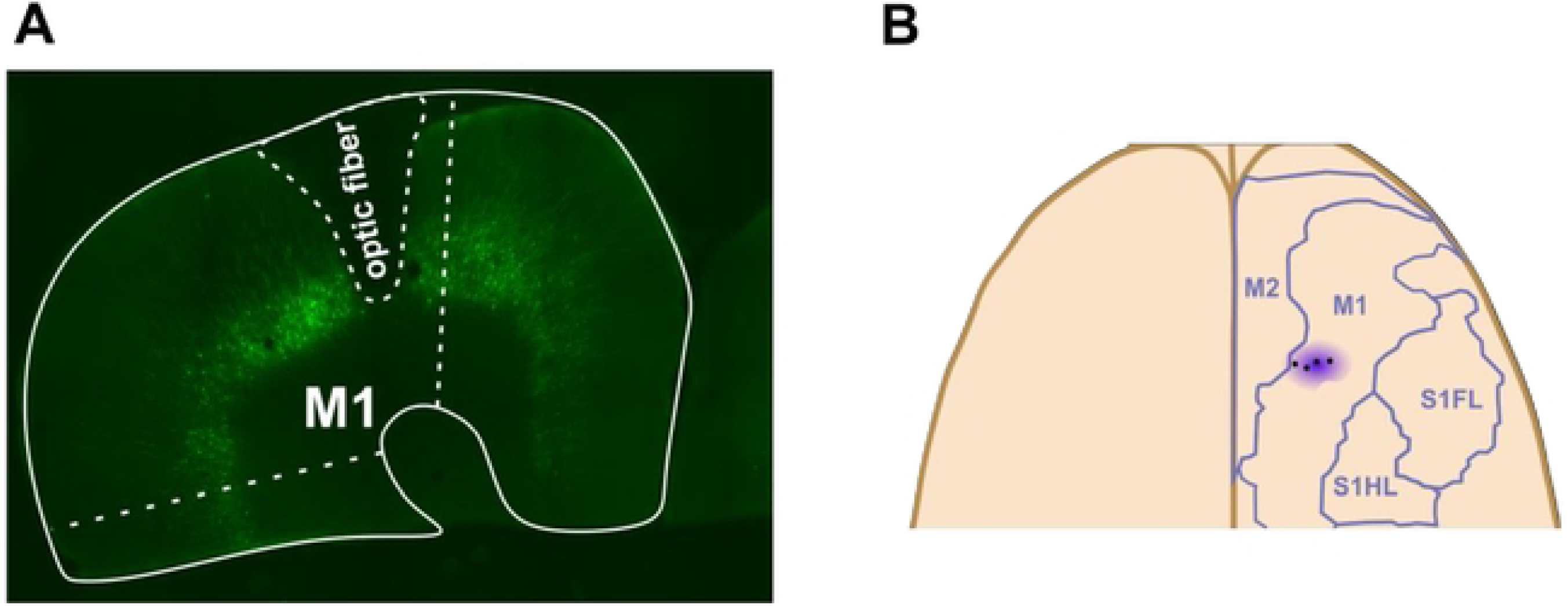
Fiber implantation in motor cortex. Coronal (A) section showing the location of the optic fiber (white dotted line). In (B), the fiber implantation area is indicated in purple shadow; the exact coordinates for each experiment are shown in black dots. SFL (anterior limb somatosensory cortex); M2 (secondary motor cortex); M1 (primary motor cortex); SHL (posterior limb somatosensory cortex)

**Supplementary Figure 4.**
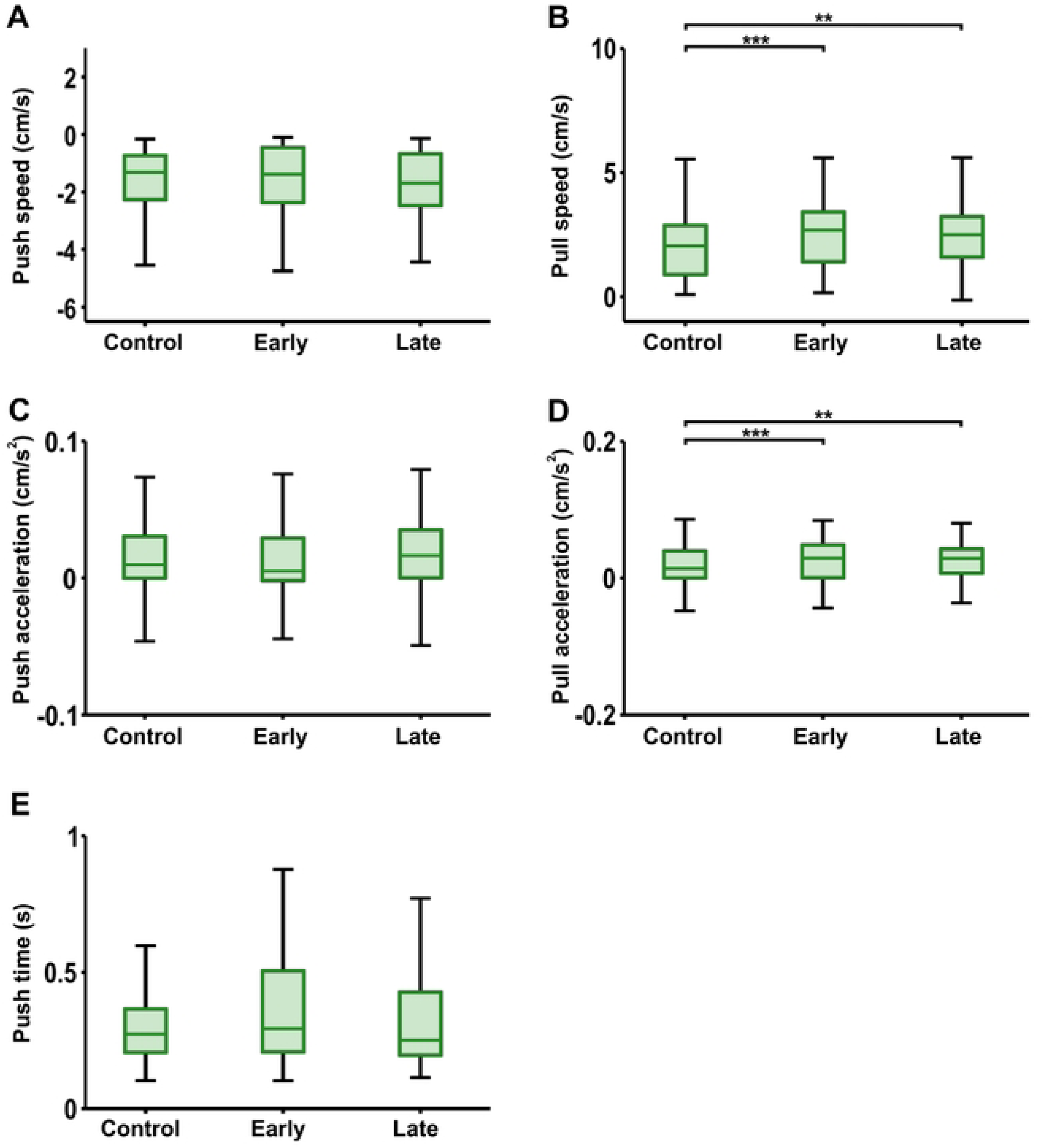
Effect of CS neuron inhibition on lever pressing trajectories. Different kinematic parameters of lever pressing movement were evaluated: push speed (A), pull speed (B), push acceleration (C), pull acceleration (D), push time (E). Comparisons were made between control lever presses vs early inhibition and control vs late inhibition (n =4; Kruskal Wallis with Dunn’s post hoc test).

**Supplementary Figure 5.**
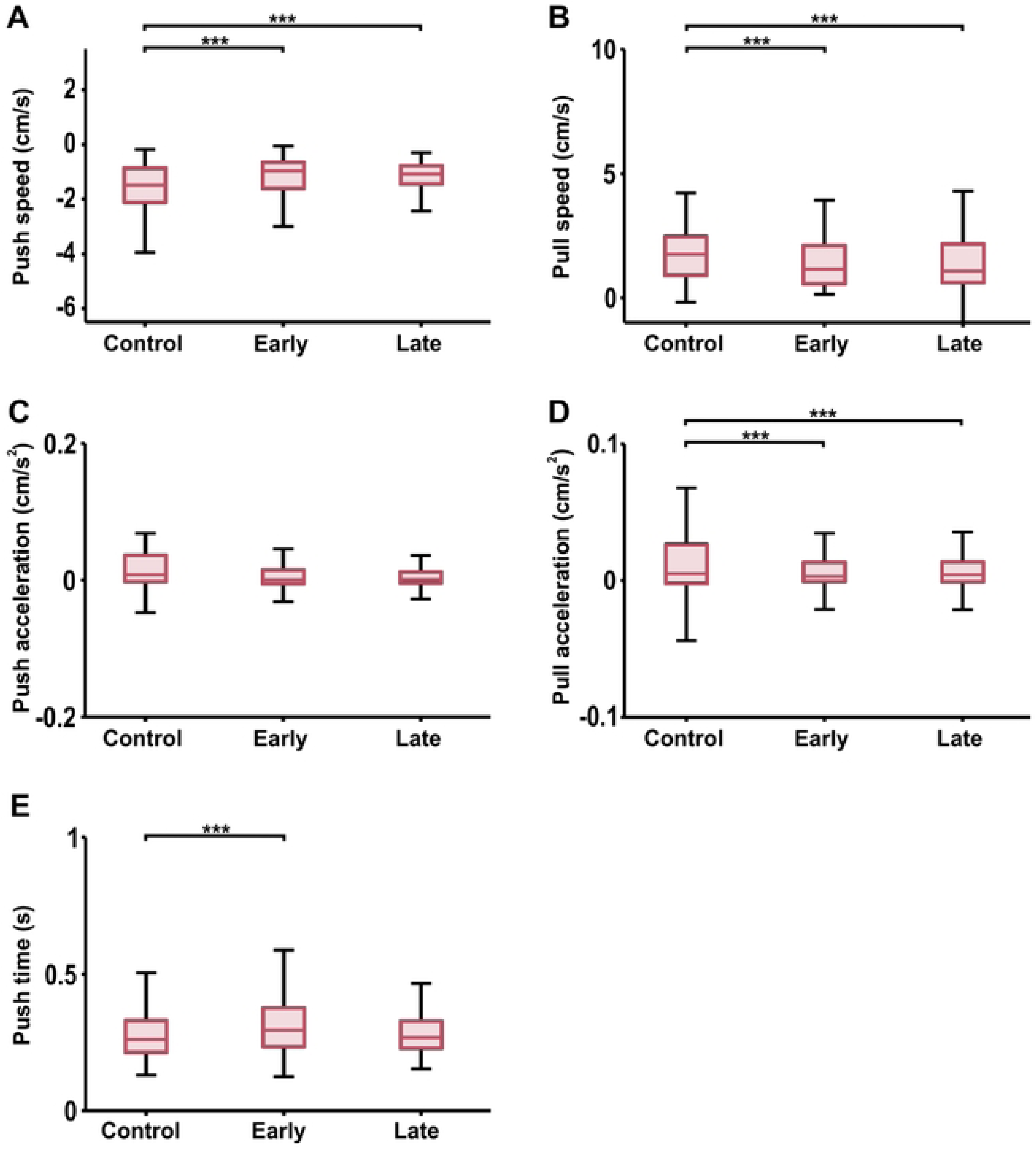
Effect of CR neuron inhibition on lever pressing trajectories. Different kinematic parameters of lever pressing movement were evaluated: push speed (A), pull speed (B), push acceleration (C), pull acceleration (D), push time (E). Comparisons were made between control lever presses vs early inhibition and control vs late inhibition (n =4; Kruskal Wallis with Dunn’s post hoc test).

